# Tryptophan derivatives regulate the seed germination and radicle growth of a root parasitic plant, *Orobanche minor*

**DOI:** 10.1101/2021.04.27.441549

**Authors:** Michio Kuruma, Taiki Suzuki, Yoshiya Seto

## Abstract

Root parasitic plant germination is induced by the host-derived chemical, strigolactone (SL). We found that a major microbial culture broth component, tryptone, inhibits the SL-inducible germination of a root parasitic plant, *Orobanche minor*. L-tryptophan (**1a**, L-Trp) was isolated as the active compound from tryptone. We further found that L-Trp related compounds (**1b**-**11**), such as a major plant hormone auxin (**8**, indole-3-acetic acid; IAA), also inhibit the germination and post-radicle growth of *O. minor*. We designed a hybrid chemical (**13**), in which IAA is attached to a part of SL, and found that this synthetic analog induced the germination of *O. minor*, and also inhibited post-radicle growth. Moreover, we found that *N*-acetyl Trp (**9**) showed germination stimulating activity, and introduction of a substitution at C-5 position incresed its activity (**12a**-**12f**). Our data, in particular, the discovery of a structurally hybrid compound that has two activities that induce spontaneous germination and inhibit subsequent radical growth, would provide new types of germination regulators for root parasitic plants.

---

Root parasitic plants in the family Orobancheceae, such as *Striga* and *Orobanche*, cause significant damage to major crops, such as sorghum, millet, maize, sugarcane and upland rice, particularly in sub-Saharan Africa. The annual loss caused by these parasitic plants is estimated to be billions of US dollars. The seed germination of these root parasitic plants is induced by host-derived chemicals collectively called strigolactones (SLs). This germination system is thought to be a survival strategy in which the parasitic plants germinate only in the host’s presence, because they cannot survive without attaching to the host. In 2005, SLs were re-characterized as symbiotic signals for arbuscular mycorrhizal fungi that aid in supplying inorganic nutrients, such as phosphate, to host plants^1^. Moreover, in 2008, it was discovered that SLs act as a novel class of plant hormones that regulate shoot branching^2, 3^.

Parasitic plants typically produce a very large number of extremely tiny (approximately 0.2 mm) seeds that may remain dormant for decades. Once the seeds detect the SL molecules emitted from host roots, they are released from dormancy, germinate and penetrate into the host roots. However, if the parasitic plant seeds germinate in the absence of a host, then they die within 5 days. A method called ‘suicidal germination’ strategy was based on this observation and is thought to be an effective way to remove the parasitic plant seeds from infected fields. However, for practical applications, large amounts of the SL molecules need to be prepared and spread on the agricultural fields. Because the amounts of naturally existing SLs are extremely small, at present, chemical synthetic approaches have been used, and SL analogs, such as the commonly used GR24, have been developed. In most cases, these synthetic analogs contain a methylbutenolide moiety (D-ring), which is a common component of natural SL molecules. The attachment of the D-ring to various structures has resulted in yielding many SL analogs^4^. Recently, the SL receptors in a parasitic plant, *Striga hermonthica*, were identified^5–7^, which greatly facilitated the screening of chemicals that target these receptor proteins. Using this approach, a femtomolar-range suicide germination inducer, sphynolactone-7, which also has the D-ring moiety, was developed recently*^8^*. In contrast, there have been limited examples of structurally SL-unrelated chemicals that have SL-like bioactivities.

In addition to these chemical synthetic or chemical biological approaches, producing SLs or chemicals with the same bioactivities in microorganisms may be useful. Some plant pathogenic microorganisms produce plant hormone molecules as secondary metabolites. For instance, *Gibberella fujikuroi*, the causal fungus of rice ‘bakanae’ disease, produces the plant hormone gibberellin (GA), and this fungus is currently used to fermentatively produce GA. GA may be used as an agrochemical, which can, in particular, generate seedless grapes.

Thus, microorganisms that can produce SLs or chemicals having SL-like bioactivities should be used to quantitatively prepare suicidal germination inducers upon root parasitic plants. Consequently, we decided to screen for microorganisms that have the ability to produce chemicals with seed germination-inducing activities for the root parasitic plant *Orobanche minor*. Before establishing an efficient screening method, we evaluated the effects of components derived from microbial culture broth on *O. minor*’s seed germination to determine whether culture broth alone affects germination. During this process, we discovered that a major culture broth component, tryptone, which is often used as a nitrogen source, strongly inhibited the GR24-inducible germination of *O. minor* seeds in a dose-dependent manner (Fig. 1A). We performed a reversed-phase high performance liquid chromatography (HPLC) analysis of tryptone and detected the inhibitory activity in a probable single-peak fraction (Fig. S1). Therefore, this fraction was collected, and the structure of the isolated chemical was determined to be a major amino acid, L-tryptophan (L-Trp) (Fig. 1B). In a previous report, the effects of 20 amino acids on the germination and radicle growth of *O. minor* seeds were determined, and several amino acids, including L-Trp, inhibited both the seed germination and radicle growth of *O. minor^9^*. Thus, our data confirmed these reported results.

**Fig. 1.**
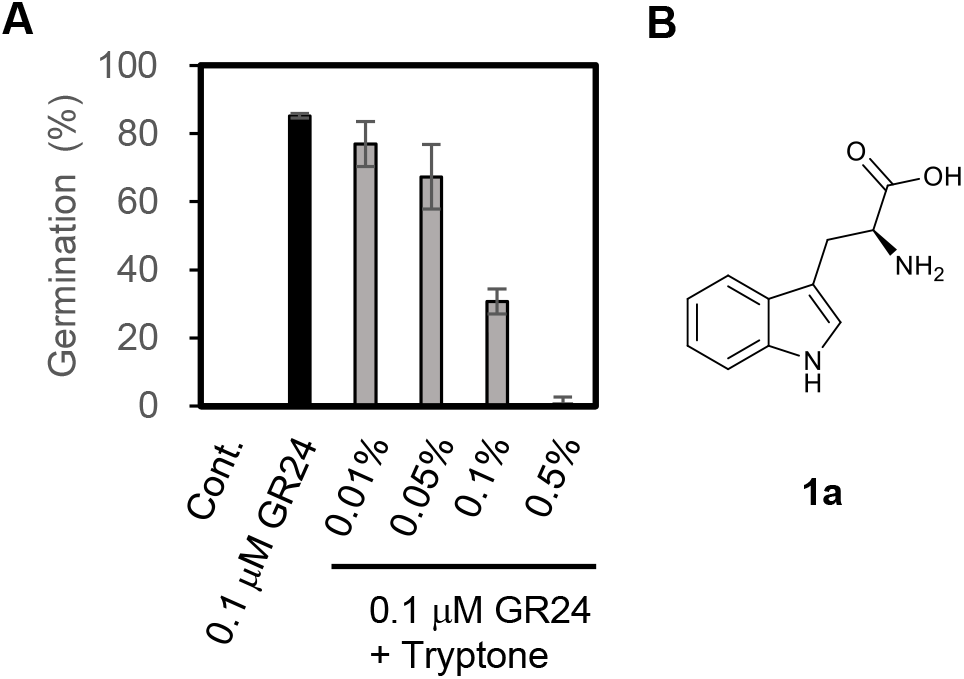
Effect of typrotone on the GR24 induced germination of *O. minor* seed. **A**. Germination rate of *O. minor* seeds after cotreatment of GR24 and different concentration of tryptone. Cont. indicates distilled water with 0.1% acetone. Data are the means ±SD (n=3). **B**. Chemical structure of isolated active compound.

To understand the structural requirements for the inhibition of germination and radicle growth by L-Trp, we tested some commercially available Trp-related chemicals. Commercial L-Trp inhibited *O. minor* germination, but the inhibitory activity of its enantiomer, D-Trp (**1b**) was relatively weak. Because some derivatives which have a substitution at C-5 position were relatively inexpensive to purchase, we tested the activity of such analogs. The introduction of a methyl group at the C-5 position (**2**) produced a weaker activity than L-Trp, whereas the introduction of a hydroxyl group at the same position (**3**) increased the activity compared to L-Trp (Fig. 2). Tryptamine (**4**) and 5-OH-tryptamine (**5**) showed weaker activity levels compared with L-Trp. L-Trp methyl ester (**6**) and L-tryptophanol (**7**) showed almost the same activity level as L-Trp (Fig. 2). *In planta*, indole-3-acetic acid (IAA; **8**), a plant hormone that is biosynthesized using Trp as a biosynthesis intermediate. Interestingly, the germination-inhibitory activity of IAA was the strongest among all the tested chemicals. In addition, L-Trp and some of the tested chemicals inhibited radicle elongation (Fig. S2)^9^, and among of them, IAA showed the strongest inhibitory effect.

**Fig. 2.**
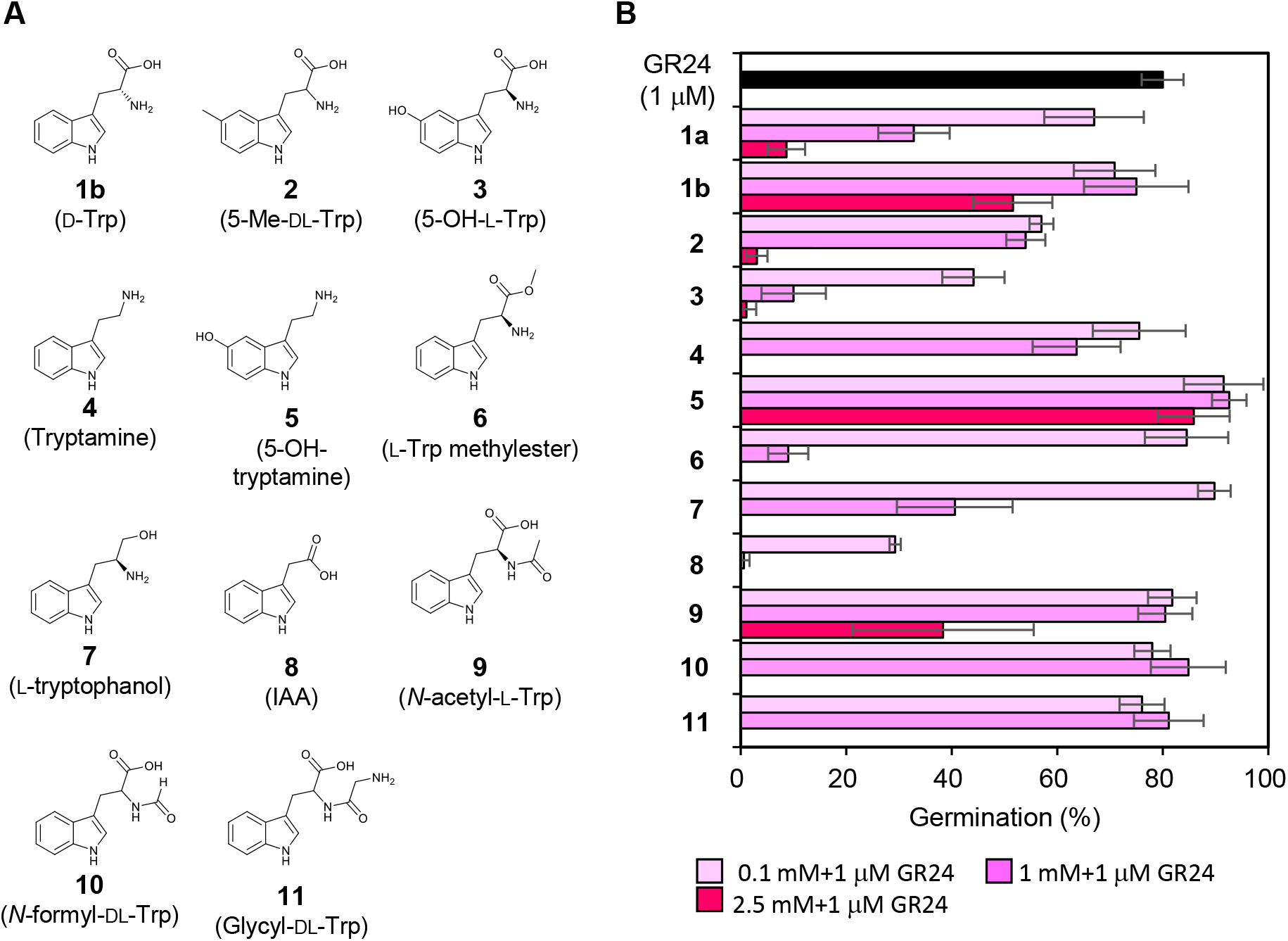
Effects of tryptophan-related compounds on *O. minor* germination. **A**.Chemical structures of tested compounds. **B**. Germination percentage of *O. minor* seed after co-treatment of GR24 (1 μM) and tested compounds at indicated concentration. Data are the means ±SD (n=4).

*N*-Acetyl-L-Trp (**9**), *N*-formyl-DL-Trp (**10**) and a dipeptide derivative, Glycyl-DL-Trp (**11**), which all have a substitution in the amino group, showed very weak germination-inhibitory activities against *O. minor* (Fig. 2B). In contrast, 2 mM *N*-acetyl-L-Trp very weakly induced the germination of *O. minor* (Fig. 3A). The activity was much weaker than that of GR24, but it was reproducible. Additionally, we chemically synthesized some derivatives of *N*-acetyl-Trp having different substitutions at the C-5 position, because a starting material, indole, which has a different substitution at C-5 position is available at a relatively low price (Fig. 3B, **12a**–**f**). Interestingly, all the C-5-substituted *N*-acetyl-Trp molecules showed stronger germination-stimulating activities than non-substituted *N*-acetyl-Trp (Fig. 3B, C). Although these activity levels were still weaker than that of GR24, these chemicals, which are not structurally related to SL, do represent new types of germination inducers for root parasitic plants.

**Fig. 3.**
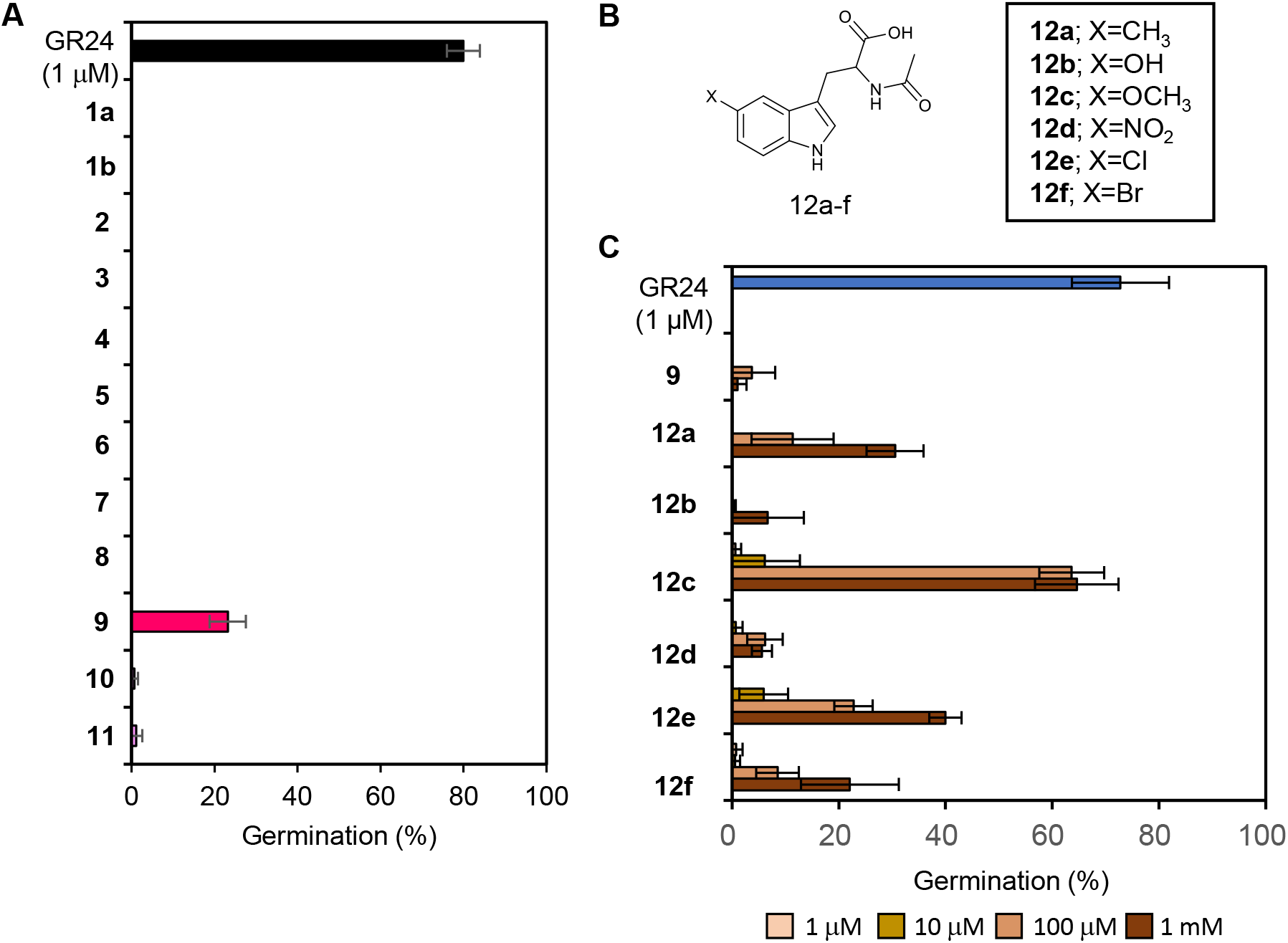
Germination stimulating activity of *N*-acetyl-tryptophan derivatives for *O. minor*. **A**. Germination stimulating activity of L-Trp related compounds. Data are the means ±SD (n=3). **B**. Chemical structures of tested compounds. **C**. Germination percentage of *O. minor* seed after treatment with tested compounds at indicated concentrations. Data are the means ± SD (n=4).

As stated above, IAA and some Trp-related chemicals inhibited not only the germination, but also the post-germination radicle growth, of *O. minor. In planta*, SLs are perceived by an α/β-fold hydrolase family protein receptor, such as DWARF14 or HYPOSENSITIVE TO LIGHT/KARRIKIN INSENSITIVE2 (HTL)^10^. In the parasitic plant *S. hermonthica*, some HTL family proteins have been identified as SL receptors required for germination^5–7^. Although the SL receptors in *O. minor* have not been identified, *Striga* HTL homologs are also conserved in *O. minor^5^*. In addition, DWARF14 and *Striga* HTL proteins possess hydrolytic activities with the SL ligand molecule^7, 11, 12^. In the SL structure, the D-ring is the most critical component for the bioactivities of SL-related chemicals. Even when a molecule that is structurally unrelated to SLs is attached to the D-ring, these synthetic analogs often show SL-like bioactivities^4^. Consequently, we chemically synthesized a hybrid SL analog in which the D-ring moiety was attached to the carboxylic acid part of IAA (Fig. 4A, **13**) to determine whether one compound might have both seed germination-inducing and radicle growth inhibitory activities. This type of hybrid chemical had been synthesized previously and had reported germination-inducing activities for several root parasitic plants, although the inhibitory activity for radical growth was not evaluated.^13^Consistent with our expectations, the synthesized compound **13** induced *O. minor* germination at moderately low concentrations and radicle growth inhibition was observed for post-germination seeds (Fig. 4B, C; Fig. S4). Thus, we could claim that compound **13** was the first case to develop a compound having the both activities. It is still unclear whether IAA was released *in planta* by receptor-dependent hydrolysis, nonspecific hydrolysis or nonenzymatic degradation. However, if this compound can suppress parasitism by inhibiting post-germination radicle growth, then such chemical can be used as a suicidal germination inducer even in the presence of host plants.

**Fig. 4.**
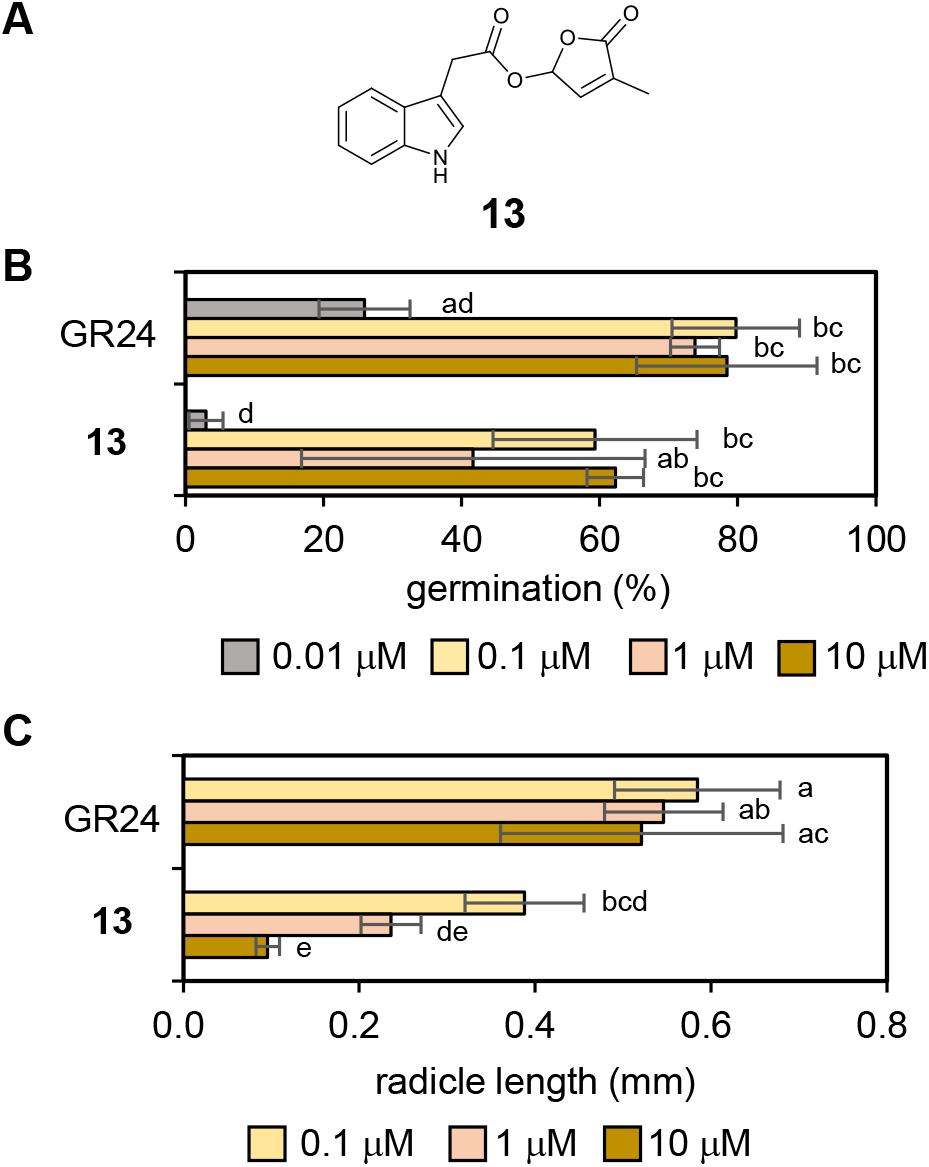
Effect of IAA-SL on germination and post-radicle growth on *O. minor*. **A**. Chemical structure of IAA-SL. **B**. Germination stimulating activity of IAA-SL. Data are the means ±SD (n=4). **C**. Radicle growth inhibitory activity of IAA-SL. Data are the means ±SD (n=5). Different letters indicate significant differences at P < 0.05 with Tukey multiple comparison test.

We identified L-Trp (**1**), which is contained in the major microbial culture broth component tryptone, as a germination inhibitor of *O. minor*. Thus, culture broth components in a bioassay-guided screening of *O. minor* seeds must be carefully selected because the components of the medium itself might exhibit inhibitory activity. Some Trp-related compounds exhibited similar germination-inhibitory activities and also inhibited radicle elongation. The activity of a plant hormone, IAA, was strongest among all the tested chemicals. In addition, it was previously reported that IAA treatments inhibit infection by the parasitic plant *Orobanche aegyptiaca^14^*. IAA is well known to regulate root growth of seed plants with inhibitory effect on main root growth at high concentration. Therefore, other tested chemicals may also be converted into IAA *in planta*, resulting in the inhibition of germination and radicle growth. On the contrary, some of the *N*-acetylated Trp derivatives showed germination-inducing activities for *O. minor*. The chemical synthesis of some analogs having substitutions at the C-5 position provided moderately active compounds; however, their activity levels were weaker than that of GR24. Most of the reported germination inducers for root parasitic plants are structural derivatives of SLs. Although the germination-stimulating activities of *N*-acetyl-Trp derivatives are weak, they are good lead chemicals for the development of new classes of suicidal germination inducers. Notably, these derivatives may be synthesized from inexpensive starting materials by a single step reaction (a substituted indole, serine or acetic anhydride), which would be an advantage for quantitative preparations. Most interestingly, we have discovered a synthetic hybrid chemical composed of IAA and the SL D-ring part, which has both germination inducing and subsequent radicle growth inhibiting activity. This is the first case to develop such SL analog. Our results provide new chemical tools to control the growth of root parasitic plants by manipulating both the germination and subsequent radicle growth steps.

## Acknowledgements

This work was funded by JSPS KAKENHI (Grant No. 19K05852), the Kato Memorial Bioscience Foundation and the Mitsubishi Foundation. We thank Dr. Xiaonan Xie for providing the *O. minor* seeds. We thank Lesley Benyon, PhD, from Edanz Group (https://en-author-services.edanz.com/) for editing a draft of this manuscript.

## Notes

### Competing Interest Statement

The authors have declared no competing interest.

