## Supplementary information for "Tryptophan derivatives regulate the seed germination and radicle growth of a root parasitic plant, *Orobanche minor*"

### Supplementary Material

Supplementary Figures

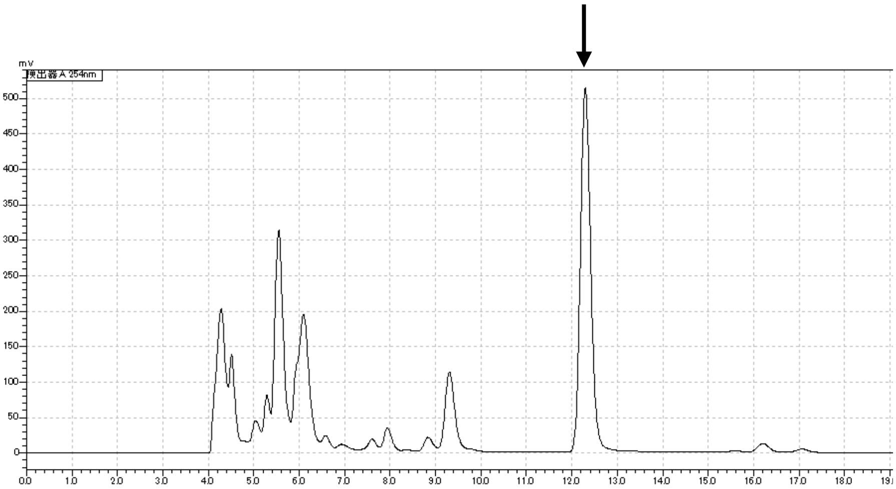


**Supplementary Figure 1.** HPLC chromatogram of tryptone detected at 254 nm. The arrow indicates the collected peak, which was determined to be $_{L}$-Trp.

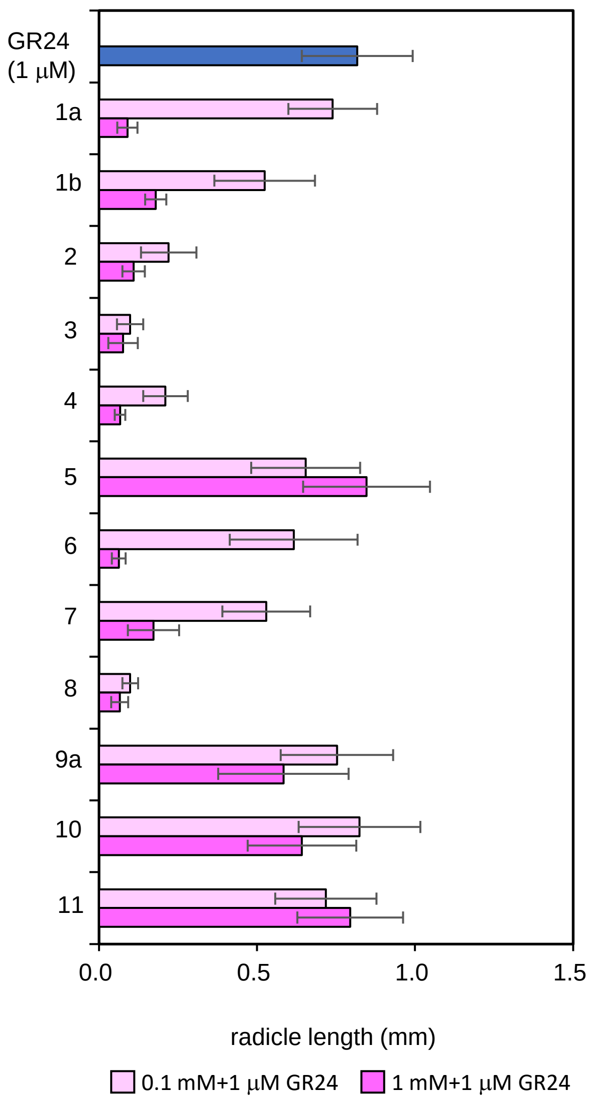


**Supplementary Figure 2.**. Effects of tryptophan-related compounds on *O. minor* radicle elongation. Radicle length of post-germinated *O. minor* after treatment with tested compounds. Data are the means ±SD (n=4-36).

(a) (0.1 mM, Scale Bar; 1 mm, GR24 0.1 μM)

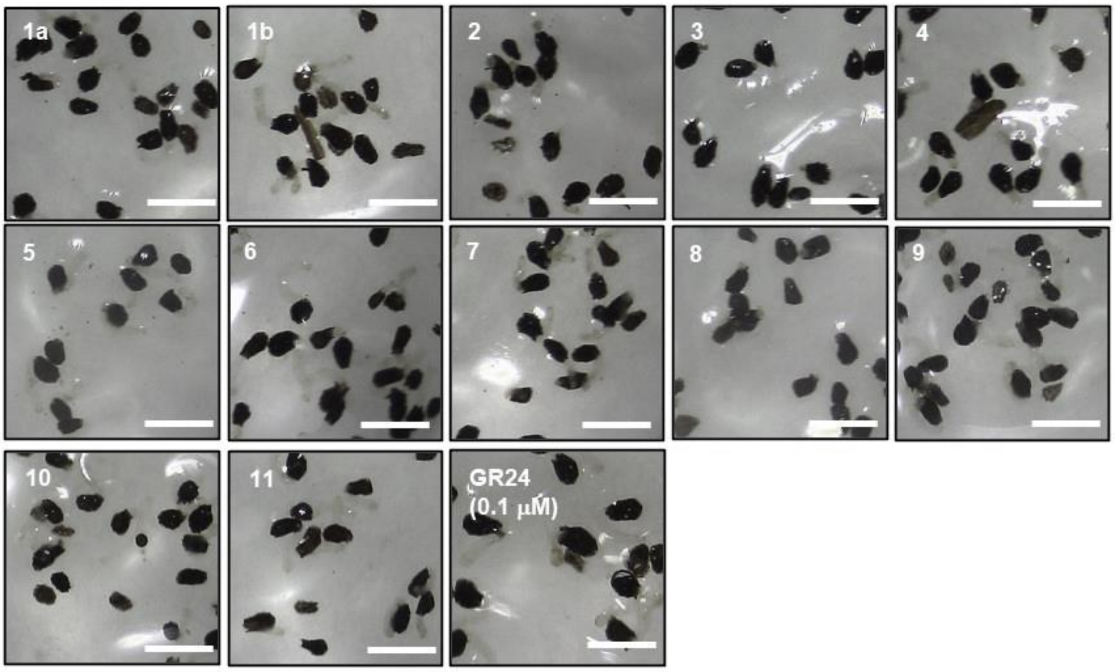


(b) (1 mM, Scale Bar; 1 mm, GR24 0.1 μM)

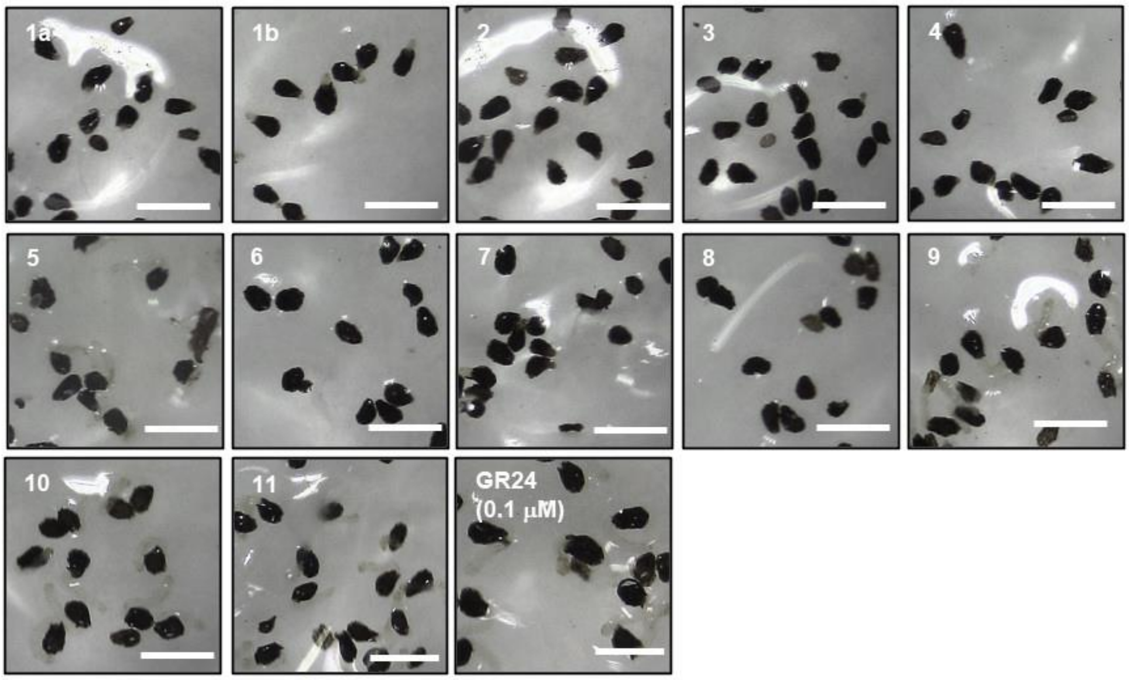


**Supplementary Figure 3.** Effects of tryptophan-related compounds on *O. minor* radicle elongation. The pictures of post-germinated radicles after the co-treatment of GR24 and each chemical at (a) 0.1 mM, and (b) 1 mM. (Scale Bar = 1 mm)

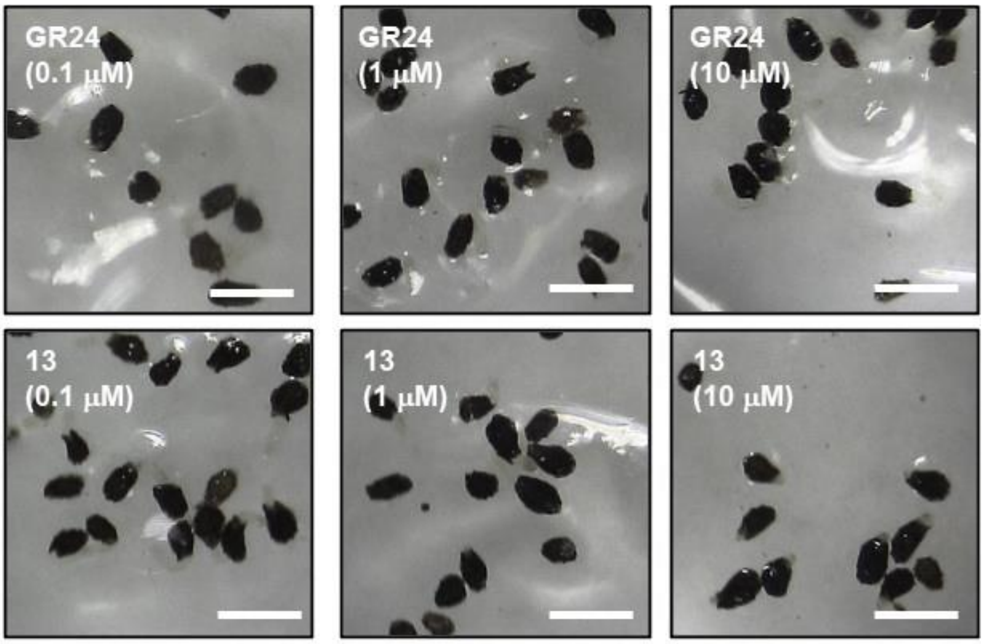


**Supplementary Figure 4.** Effects of IAA-SL (13) on O. minor radicle elongation. The pictures of post-germinated radicles after the treatment of GR24 or IAA-SL at the indicated concentration (Scale Bar = 1 mm)

Tryptophan derivatives regulate seed germination and radicle growth of a root parasitic

### Materials and Methods

#### Generals

HR-MS was recorded with LC-Q-Tof-MS (SCIEX X500R), equipped with a reverse phase HPLC separation (Acquity UPLC BEH-C18, ϕ2.1 × 50 mm, 1.7 μM)). NMR spectra were recorded with JEOL AL300 ($^1$H at 300 MHz and $^{13}$C at 75 MHz).

#### Chemicals

L-Trp, L-serine, and 5-OH-tryptamine were purchased from FUJIFILM Wako Pure Chemical Corporation (Osaka, Japan). 5-hydroxy-DL-tryptophan were purchased Alfa Aesar (Massachusetts, United States). Other chemicals were purchased by TOKYO CHEMICAL INDUSTRY CO., LTD. (Tokyo, Japan). Other chemicals were synthesized as described below.

#### Isolation of a germination inhibitor from tryptone

Tryptone was dissolved with water, and subjected to a reverse phase HPLC (Mightysil, RP18, $\phi$4.6x250 mm, 20% MeOH/$H_2O$, 0.6mL/min). A peak at 12.3 min (Fig. S1) was collected to give a compound (6.4 mg). $^1$H-NMR (300 MHz, $CD_3OD$) $\delta$ 7.63 (d, $J$ = 7.5 Hz, 1 H) 7.29 (d, $J$ = 7.5 Hz, 1H) 7.12 (s, 1H) 7.05 (t, $J$ = 7.0 Hz, 1H) 6.97 (t, $J$ = 7.0 Hz, 1H) 3.78 (dd, $J$ = 9.6, 3.9 Hz, 1H) 3.45 (dd, $J$ = 15, 3.9 Hz, 1H) 3.07 (dd, $J$ = 15, 9.4Hz, 1H)

#### Determination of stereochemistry of the isolated tryptophan

The stereochemistry of isolated Trp was determined using a modified Marfey's method. The isolated Trp, and the standard L-Trp, and L-Trp, were separately dissolved in water (1 mg/mL). A 100 μL of FDLA solution (1% in acetone) and 20 μL of 1N $NaHCO_3$ were added to each amino acid solution (50 μL), and incubated at 45°C for 1 h. Twenty μL of 1 N HCl was added to each reaction solution, and the samples were diluted by adding 810 μL of acetonitrile. 10 μL of each sample was analysed by a reverse phase HPLC (Mightysil, RP18, $\phi$4.6x250 mm, 60% acetonitrile/0.1%TFA water, 0.6mL/min).

The derivative from standard L-Trp and D-Trp were detected at 16.0 min and 22.7min, respectively. The derivative from the isolated Trp was detected at 16.0 min.

**Chemical synthesis of *N*-acetyl-tryptophan derivatives**

Preparation of *N*-acetyl-5-Methoxy-DL-Trp (**12c**), *N*-acetyl-5-chloro-DL-Trp (**12e**), *N*-acetyl-5-bromo-DL-Trp (**12f**), and *N*-acetyl-5-nitro-DL-Trp (**12d**).

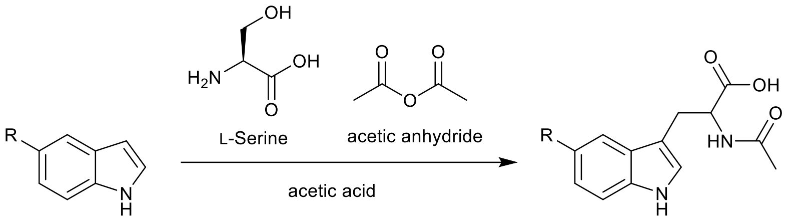


Chemical synthesis of *N*-acetyl-5-Methoxy-DL-Trp, *N*-acetyl-5-chloro-DL-Trp, *N*-acetyl-5-bromo-DL-Trp, and *N*-acetyl-5-nitro-DL-Trp was performed according to a reported method[1]. L-serine (2 mmol) was added to a solution of the substituted indole (1 mmol) and acetic anhydride (9 mmol) in acetic acid (2.4 ml). In the case of *N*-acetyl-5-nitro-DL-Trp, 2.5 mmol L-serine was used. The mixture was stirred under argon at 73°C for 2 h. Upon cooling to room temperature, the solution was adjusted to 30 ml with water. Water layer was extracted with EtOAc (3×30 ml). The combined organic extracts were dried over $Na_2SO_4$, and then concentrated *in vacuo*, and purified by a silica gel column chromatography ($CHCl_3$/MeOH 9:1) to afford *N*-acetyl-DL-Trp derivatives.

Preparation of *N*-acetyl-5-methyl-DL-Trp (**12a**) and *N*-acetyl-5-hydroxy-DL-Trp (**12b**).

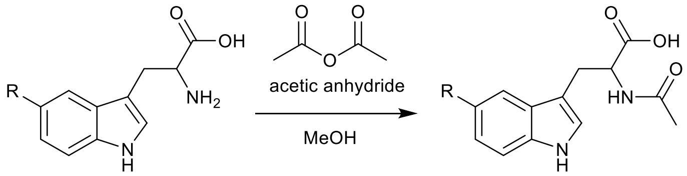


Chemical synthesis of *N*-acetyl-5-methyl-DL-Trp and *N*-acetyl-5-hydroxy-DL-Trp were performed according to a reported method[2]. 5-methyl-DL-Trp (0.2 mmol) was added to a solution of acetic anhydride (0.8 mmol) in MeOH (1 ml). 5-hydroxy-DL-Trp (1.0 mmol) was added to a solution of acetic anhydride (6.0 mmol) in MeOH (15 ml). The mixture was stirred at 40°C for 16 h, and the solution was concentrated *in vacuo*. The crude mixture was purified by a silica gel column chromatography (*n*-hexane/EtOAc 2/8) to afford *N*-acetyl-5-methyl-DL-Trp. 5-OH-L-Trp (0.4 mmol) was added to a solution of acetic anhydride (0.8 mmol) in MeOH (2 ml). The mixture was stirred at 40°C for 16 h and the solution was concentrated *in vacuo*. The crude mixture was purified by a silica gel column chromatography (*n*-hexane/EtOAc:1/9) to afford *N*-acetyl-5-OH-L-Trp.

**Preparation of IAA-SL (13)**

$K_2CO_3$ (1.0 mmol) was added to the solution of IAA (0.50 mmol) in 5.0 mL of *N*-methyl-2-pyrrolidone. 5-bromo-3-methyl-2(5H)-franone (0.98 mmol) was added to the mixture, and the mixture was stirred for 12 h at room temperature. The reaction was quenched by adding 1 N HCl, and the solution was diluted to 50 mL with water, extracted with EtOAc (3×50 mL). The combined organic layers were dried over $Na_2SO_4$, and then concentrated *in vacuo*. The crude sample was purified by a silica gel column chromatography (n-hexane/EtOAc:6/4) to give IAA-SL (95 mg, 70% yield) as yellow oil.

**Spectral data of synthetic chemicals**

*N*-acetyl-5-Methoxy-DL-Trp (**12c**)

Yield: 144 mg (52%). $^1$HNMR (300 MHz, $CD_3OD$) $\delta$ 7.20 (d, *J* = 9.0 Hz, 1 H),7.05 (d, *J* = 9.0 Hz, 2 H), 6.74 (dd, *J* =8.7, 2.4 Hz, 1 H), 4.71 (dd, *J* = 7.8, 5.1 Hz, 1 H), 3.33-3.26 (m, 1 H), 3.13 (dd, *J* = 11.9, 8.3 Hz, 1 H), 1.91 (s, 3H), HRMS [ESI- (*m*/*z*)] calculated for $(C_{14}H_{16}N_2O_4\ \text{-H})^-$ 275.1037, found 275.1050.

*N*-acetyl-5-nitro-DL-Trp (**12d**)

Yield: 18 mg (6%). $^1$HNMR (300 MHz, $CD_3OD$), $\delta$ 8.58 (d, *J* = 2.4 Hz, 1 H), 8.02 (dd, *J* = 9.3, 2.1 Hz, 1 H), 7.43 (d, *J* =9.0 Hz, 1 H), 7.31 (s, 1 H), 4.75 (dd, *J* = 7,5, 5.1 Hz, 1 H), 3.33-3.26 (m, 1 H), 3.22 (dd, *J* = 14.7, 7.8 Hz, 1 H), 1.92 (s, 3H), HRMS [ESI- (*m*/*z*)] calculated for $(C_{13}H_{13}N_3O_5\ \text{-H})^-$ 290.0782, found 290.0797

*N*-acetyl-5-chloro-DL-Trp (**12e**)

Yield: 73 mg (26%).$^1$HNMR (300 MHz, $CD_3OD$) $\delta$ 7.53 (d, *J* = 1.8 Hz, 1 H), 7.28 (d, *J* = 8.4 Hz, 1 H), 7.14 (s, I H), 7.04 (dd, *J* =8.7, 2.1 Hz, 1 H), 4.69 (dd, *J* = 7.7, 5.3 Hz, 1 H), 3.33-3.26 (m, 1 H), 3.13 (dd, *J* = 11.9, 8.3 Hz, 1 H), 1.91 (s, 3H), HRMS [ESI- (*m*/*z*)] calculated for ($C_{13}H_{13}ClN_2O_3$ -H)$^-$ 279.0542, found 279.0558.

*N*-acetyl-5-bromo-DL-Trp (**12f**)

Yield: 285 mg (88%).$^1$HNMR (300 MHz, $CD_3OD$) $\delta$ 7.69 (d, *J* = 1.2 Hz, 1 H), 7.24 (d, *J* = 7.8 Hz, 1 H), 7.16 (dd, *J* = 8.7, 1.8 Hz, 1H), 7.12 (s, 1 H), 4.67 (dd, *J* = 7.5, 5.1 Hz, 1 H), 3.32-3.27 (m, 1 H), 3.11 (dd, *J* = 14.7, 7.8 Hz, 1 H), 1.91 (s, 3H), HRMS [ESI- (*m*/*z*)] calculated for ($C_{13}H_{13}BrN_2O_3$ -H)$^-$ 323.0037, found 323.0057.

*N*-acetyl-5-methyl-DL-Trp (**12a**)

Yield: 15 mg (29%). $^1$HNMR (300 MHz, $CD_3OD$), $\delta$ 7.34 (s, 1 H), 7.19 (d, *J* = 8.4 Hz,1 H), 7.03 (s, 1H), 6.91 (dd, *J* = 7.1, 1.1 Hz), 4.69 (dd, *J* = 7.7, 5.0 Hz, 1 H), 3.32-3.25 (m, 1 H), 3.07 (dd, *J* = 14.7, 8.0 Hz, 1 H), 2.41 (s, 3H), 1.90 (s, 3H),$^{13}$C NMR (300 MHz, $CD_3OD$), $\delta$ 21.72, 22.45, 28.54, 54.99, 110.61, 111.99, 118.91, 124.03, 124.43, 128.84, 129.16, 136.43, 173.20, 175.56, HRMS [ESI- (*m*/*z*)] calculated for ($C_{14}H_{16}N_2O_3$ -H)$^-$ 259.1088, found 259.1094.

*N*-acetyl-5-hydroxy-DL-Trp (**12b**)

Yield: 67 mg (26%). $^1$HNMR (300 MHz, $CD_3OD$), $\delta$ 7.15 (d, *J* = 8.4 Hz, 1 H),7.02 (s, 1 H), 6.94 (d, *J* =2.3 Hz, 1 H), 6.66 (dd, *J* = 8.6, 2.3 Hz, 1 H), 4.69 (dd, 8.1, 5.1 Hz, 1 H), 3.32-3.26 (m, 1 H), 3.13 (dd, *J* = 14.6, 8.1 Hz, 1 H), 1.91 (s, 3H), $^{13}$CNMR (300 MHz, $CD_3OD$), $\delta$ 22.32, 28.44, 54.66, 103.34, 110.19, 112.38, 112.63, 125.00, 129.47, 132.85, 151.20, 173.16, 175.30, HRMS [ESI- (*m*/*z*)] calculated for ($C_{13}H_{14}N_2O_4$ -H)$^-$ 261.0891, found 261.0889.

IAA-SL (**13**)

Yield: 94.9 mg (70%). $^1$HNMR (300 MHz, $CDCl_3$) $\delta$ 8.19 (brs, 1 H), 7.58 (d, *J* = 7.8 Hz, 1 H), 7.37(d, *J* =7.5 Hz, 1 H), 7.26-7.12 (m, 3 H), 6.89-6.85 (m, 2 H), 3.86 (d, *J* = 1.2 Hz, 2 H), 1.96 (t, *J* =1.7 Hz, 3 H)

### *Orobanche minor* germination assay

*Orobanche minor* seeds were washed with 70% EtOH, and then sonicated for 4 min in 1% sodium hypochlorite solution containing 0.2% Tween-20. The seeds were then washed ten times with sterile water and suspended in 0.1% agar solution. The seeds were loaded onto 5-mm glass fiber filter disks (20-70 seeds/disk), and were conditioned at 23$^{o}$C for 15 days. Each disk was transferred into a 96-well plate. A 30 µl aliquot of test chemical solution was added to the well. For the germination assay, the chemical solutions were prepared by 1000 times dilution from each acetone stock solution with water (final acetone concentration was 0.1%). GR24 solution (0.1% acetone) and sterile water (0.1% acetone) was used as positive and negative control, respectively. The 96 well plates were incubated for 5 days at 23$^{o}$C. For germination inhibitory assays, 1 or 0.1 mM GR24 solution in acetone stock was 1000 times diluted with a test chemical solution in water (final GR24 concentration was 1 or 0.1 µM). The concentration of test chemical was determined due to the solubility of each chemical (Tryptamine, indole-3-acetic acid, $N^{\alpha}$-formyl-Trp, and *N*-glycyl-Trp were tested at a maximum concentration of 1 mM, and other chemicals were tested at a maximum concentration of 2.5 mM). Radicle length of germinated seeds were measured using image J (https://imagej.nih.gov/ij/). For inhibitory assay using tryptone, 0.1 mM GR24 in acetone stock was 1000 times diluted with a tryptone solution in water at indicated concentration (0.01%, 0.05%, 0.1%, 0.5%).
